# Early-life adversity mediates a thalamo-amgydalar circuit dysfunction underlying chronic pain and anxiety

**DOI:** 10.1101/2024.12.13.628438

**Authors:** Anna Maria Borruto, Tabea Kohlund, Martin von Renesse, Lukas Rettig, Nikolas Leonhardt, Claudia Calpe-López, Luca Benn, Hans-Christoph Friederich, Rohini Kuner, Thomas Kuner, Rainer Spanagel, Sebastian Wieland

**Author notes:** <u>Contact information/Corresponding author:</u> Sebastian Wieland; Department for General Internal Medicine and Psychosomatics; University Hospital Heidelberg; Thibautstr. 4; 69115 Heidelberg; 06221-5634180. Equal contribution (Anna Maria Borruto, Tabea Kohlund, Martin von Renesse).

## Abstract

Childhood adversity increases the risk of developing a vicious cycle of chronic pain and comorbid anxious avoidance, yet the underlying biological mechanisms remain unclear. Here, we investigated the role of a brain circuit from the paraventricular thalamus (PVT) to the central amygdala (CeA) in mediating hyperalgesia and comorbid anxiety following early life stress. Using a vulnerability-stress model, we exposed both male and female mice to early social isolation (vulnerability) followed by nerve injury (stress) and showed increased hyperalgesia and anxious avoidance behavior in nerve-injured female mice following early adversity. Chemogenetic, electrophysiological, and optophysiological analyses revealed a causal contribution of a hyperexcitable PVT-CeA circuit dysfunction to chronic hyperalgesia and anxiety in nerve-injured female mice after early life stress. Our findings reveal a neural mechanism linking childhood adversity to chronic pain and anxiety, and suggest that reprogramming this pathway may reverse the impact of childhood adversity.

## Introduction

Comorbid anxiety disorders in chronic pain patients represent a significant medical challenge reducing the efficacy of standard therapies due to increased avoidance behavior, heightened fear of pain and emotional distress^1,2^. Mental disorders like anxiety disorders develop as the result of an interaction between a predisposition (vulnerability) and a triggering event, such as a stressful life experience^3^. Early-life adversity, including childhood abuse and deprivation, represents a key vulnerability factor for the development of chronic pain^4^ and anxiety disorders^5^, in particular in women^6^. Although the type and frequency of childhood adversity partly explain the increased prevalence observed in women, emerging evidence points to additional biological mechanisms specific to women^7^. Disentangling the causal contributions of childhood adversity and biological factors remains a significant challenge in clinical research due to variability in early life stressors, the long temporal gap between childhood and adulthood, and limited control over daily stress exposure and causal analyses.

Rodent studies have shown that early adversity increases anxiety and hyperalgesia with variable sex-dependent effects^8–11^, but the underlying circuit mechanisms remain unknown. Both early adversities and acute stressors^12,13^ activate the paraventricular thalamic nucleus (PVT), a brain region that dynamically encodes stimulus salience^14^ and gates adaptive behavioral responses, particularly under conditions of motivational conflict^15^. Switching between opposing fear responses is mediated, inter alia, by PVT projections to the central amygdala (CeA)^16^. CeA-projecting PVT neurons are also involved in the maintenance and retrieval of long-term fear memories^12,16,17^ and contribute to pain processing^18–20^. Since anticipation of threat or pain^21^ also activates the midline thalamus and CeA in humans^22^, the PVT-to-CeA pathway is considered a key circuit for threat processing^16,23–25^. Current hypotheses suggest that childhood adversity alters threat processing in the PVT-to-CeA pathway that drives a trajectory towards psychopathology^26^. In this study, we now test the hypothesis that early adversity induces PVT→CeA circuit dysfunction that drives a trajectory towards heightened hyperalgesia and comorbid anxiety after a nerve lesion, especially in female mice.

## Results

### Vulnerability-stress conditions induce comorbid hyperalgesia and anxiety in female mice

To explore how early adversity affects the development of chronic pain and comorbid anxiety after a nerve lesion, we established a vulnerability-stress model in male and female mice. Early social isolation after weaning (SI; from postnatal day 21) served as a vulnerability factor, while spared nerve injury (SNI) at weeks 5-8 served as a somatic injury and additional stressor. Long-term behavioural changes in the chronic pain state were assessed in SI+SNI mice of both sexes over a period of 3 months. Control mice (ctrl) were housed in groups and underwent sham surgery. To assess the specificity of these effects, additional groups included mice subjected to social isolation only (SI-only) or spared nerve injury only (SNI-only). The experimental timeline and groups are shown in **Fig. 1A** and **1B**, respectively.

**Figure 1.**
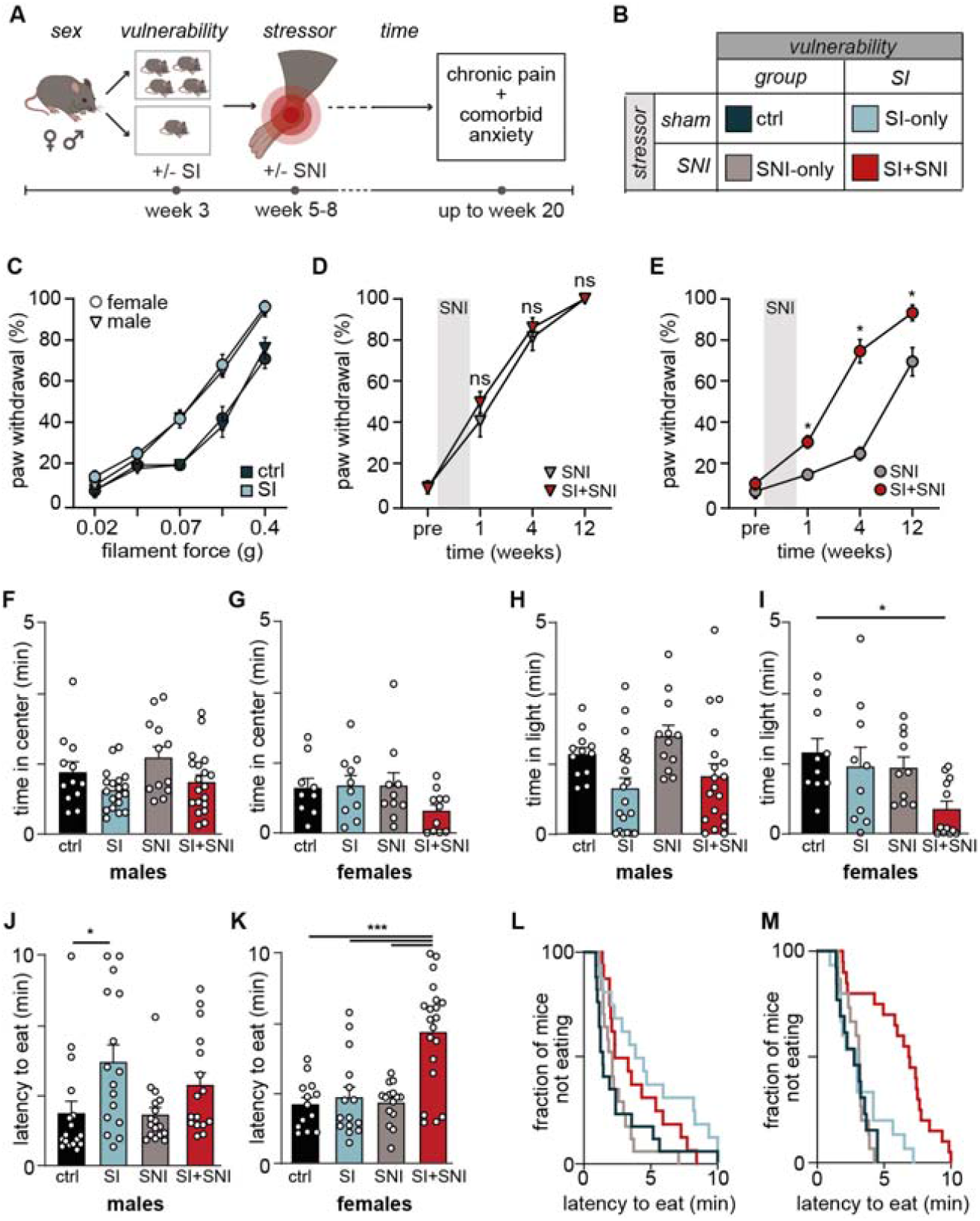
Vulnerability-stress conditions induce chronic hyperalgesia and anxious avoidance in female mice. **A.** Schematic illustration of the vulnerability-stress model. SI, social isolation. SNI, spared nerve injury. **B**. Experimental groups with corresponding color coding. **C**. Changes in withdrawal responsivity to filament force during von Frey test in male and female control (ctrl) and SI-only mice 12 weeks after sham surgery; male ctrl (n=12), SI-only (n=19) mice; F_12,174_=11.25, p(force x SI)<0.0001; female ctrl (n=10), SI-only (n=10) mice; F_12,128_=4.15, p(force x SI)<0.0001). Two-way ANOVA followed by Tukey’s post-hoc test. **D-E**. Longitudinal effects of SI on mechani-cal hypersensitivity to a 0.02 g von Frey filament at 1, 4, and 12 weeks after SNI surgery in males (**D**) and females (**E**); male SNI-only (n=12), SI+SNI (n=19); F_3,87_=0.59, p(force x SI)=0.620); female SNI-only (n=10), SI+SNI (n=12); F_3,60_=10.34, p(force x SI)<0.0001. Two-way ANOVA followed by Tukey’s post-hoc test. F-G. The time spent in the center zone during the OFT in (**F**) male ctrl (n=12), SI-only (n=19), SNI-only (n=12), SI+SNI (n=19) mice; F_1,58_=0.175; p(SIxSNI)=0.677; p(SI)=0.011 and in (**G**) female ctrl (n=9), SI-only (n=10), SNI-only (n=10), SI+SNI (n=11) mice; F_1,36_=1.45; p(SIxSNI)=0.236; p(SI)=0.202. Two-way ANOVA followed by Tukey’s post-hoc test. **H-I**. Time spent in the light compartment during the Light/Dark Box test in (**H**) male ctrl (n=12), SI-only (n=19), SNI-only (n=12), SI+SNI (n=18) mice; F_1,58_=0.067; p(SIxSNI)=0.796; p(SI)=0.002 and in (**I**) female ctrl (n=10), SI-only (n=10), SNI-only, (n=10), SI+SNI (n=12) mice; F_1,38_=1.01; p(SIxSNI)=0.322; p(SI)=0.046; ^*^p=0.021. Two-way ANOVA followed by Tukey’s post-hoc test. **J-M**. Avoidance behavior measured as latency to eat during the NSF test in (**J**) male ctrl (n=17), SI-only (n=16), SNI-only (n=17), SI+SNI (n=16) mice; X^2^=10.62, DF=3, p=0.014 and in (**K**) female ctrl (n=13), SI-only (n=15), SNI- only (n=15), SI+SNI (n=20) mice; X^2^=27.71, DF=3, p<0.0001 showing that the interaction of SI and SNI increases avoidance behavior in female mice. Log-rank (Mantel-Cox) with Bonferroni post-hoc correction. **L-M**. Corresponding survival curves of male (**L**) and female (M) mice during the NSF test, depicting the fraction of mice that did not eat during the test.

To investigate whether SI induces chronic sex-specific hyperalgesia in adult mice, we performed the von Frey test before and 12 weeks after sham surgery. We observed that SI increased paw withdrawal sensitivity similarly in both male and female mice (**Fig. 1C**) suggesting pronociceptive effects of social isolation. As early adversity may serve as a predisposition for the chronification of neuropathic pain in adult mice, we performed the von Frey test before and at 1, 4 and 12 weeks after SNI surgery and found that SI specifically increased paw withdrawal sensitivity in nerve-injured female (but not male) mice (**Fig. 1D-E**). In the cold plate test, SI did not alter thermal sensitivity in male and female mice (**Fig S2**). We conclude that SI acts as a general stressor that increases mechanical sensitivity in both sexes, while also acting as a sex-specific modulator that longitudinally enhances neuropathic pain in female mice, without affecting thermal sensitivity in either sex.

To determine whether a vulnerability-stress interaction also contributes to anxious avoidance in a sex-specific manner, we conducted a battery of anxiety tests in male and female mice at least six weeks after SNI (or sham) surgery. These tests included the Open Field Test (OFT), the Light-Dark Box Test (LDB) and the Novelty-Suppressed Feeding Test (NSF) which assesses conflict anxiety by requiring the animal to withstand an anxiogenic environment in order to obtain a reward^27^. All tests depend on active exploration and neither SI nor SNI significantly reduced locomotor activity in male and female mice (**Fig. S2**). In male mice, SI increased avoidance behavior in the OFT (**Fig. 1F**), LDB (**Fig. 1H**) and NSF (**Fig. 1J**) without an interactional effect of SNI suggesting that SI elevates anxiety levels in male mice without an additive effect of vulnerability-stress conditions (**Fig. 1J and 1L**). In female mice, neither SI nor SNI significantly affected exploratory behavior in the OFT, whereas SI and SNI induced significant avoidance of the light chamber specifically in female SI+SNI mice in the LDB (**Fig. 1H-I**). Remarkably, the interaction of SI and SNI strongly increased anxious avoidance under motivational conflict during NSF in female SI+SNI mice (**Fig. 1K and 1M**). This effect was absent in all other female groups (ctrl, SI-only, and SNI-only; **Fig. 1K and 1M**). In parallel, the latency to eat and overall food consumption in the home cage remained unchanged in male and female mice (**Fig. S3**) suggesting that the exposure to a novel anxiogenic environment mediates the anxious avoidance in the NSF test. We also observed that most (but not all) female SI+SNI mice developed strong anxious avoidance behavior (**Fig. 1M**). These results demonstrate that vulnerability-stress interaction induces a sex-specific increase in conflict anxiety and light avoidance in female SI+SNI mice, whereas male mice do not show additive avoidance behavior upon vulnerability-stress conditions. Variability among female SI+SNI mice suggests individual differences in resilience to vulnerability-stress conditions.

### Vulnerability-stress conditions induce presynaptic facilitation and hyperexcitability of the PVT→CeA pathway in female mice

In order to virally target CeA-projecting PVT neurons, we stereotaxically injected mCherry-expressing adeno-associated virus (AAVretro-mCherry) in the CeA and detected retrograde mCherry expression in PVT neurons (**Fig. 2A**). To characterize the synaptic connection of the PVT→CeA pathway, we injected Channelrhodopsin-2-expressing AAV (AAV-ChR2) in the PVT and prepared acute CeA slices containing ChR2-expressing PVT terminals (**Fig. 2B**). Brief light pulses evoked robust α-amino-3-hydroxy-5-methyl-4-isoxazolepropionic acid receptor (AMPAR) and N-methyl-d-aspartate receptor (NMDAR)-mediated excitatory postsynaptic currents (EPSCs) (**Fig. 2C**) in 284 of 322 recorded CeA neurons (88.2%), as they were pharmacologically blocked by AMPAR antagonist CNQX and NMDAR antagonist AP-V (**Fig.S4**). Light stimulation of PVT terminals evoked EPSCs with a short delay of 5.55±0.07 ms suggesting that glutamatergic PVT terminals directly innervate CeA neurons.

**Figure 2.**
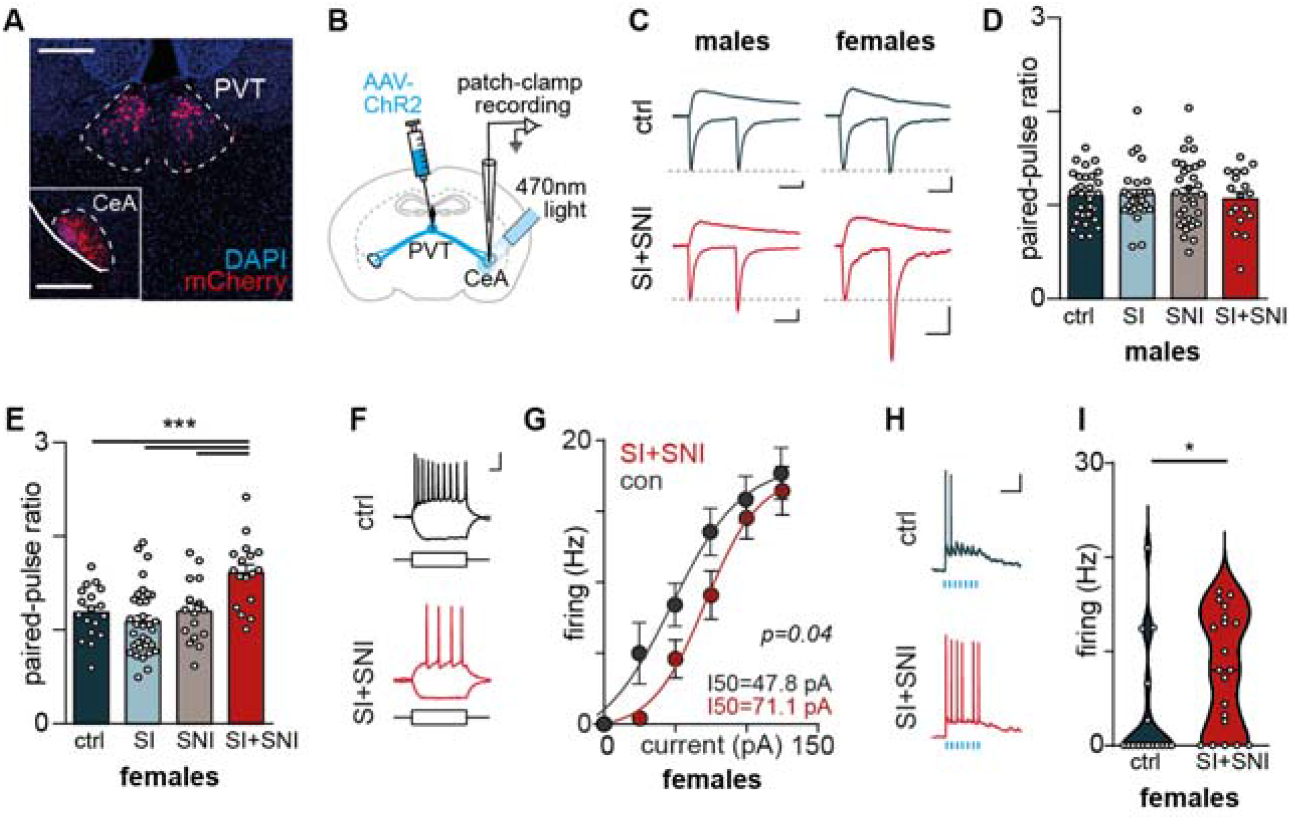
Vulnerability-stress conditions induce presynaptic plasticity and hyperexcitability of the PVT→CeA pathway in female mice **A.** Retrogradely labeled PVT neurons were observed 3 weeks after injection of AAVretro-mCherry into the CeA. Scale bar=300 μm, inset shows the injection site. **B**. Schematic optogenetic stimulation of PVT axons during in- vitro recordings in CeA neurons. **C**. Example traces of light-evoked EPSCs at ™70 mV and +40 mV, Scale bar=50 pA, 50ms. **D-E**. Paired-pulse ratio in (**D**) male ctrl (n=30 cells), SI-Only (n=27), SNI-Only (n=32), SI+SNI (n=18) F_3,103_=1.383, p=0.932 and (**E**) female ctrl (n=18 cells), SI-Only (n=33), SNI-Only (n=16), SI+SNI (n=17), F_1,80_=11.02, p(SIxSNI)=0.0014, F_3,80_= 0.393, p<0.0001. Two-Way ANOVA followed by post-hoc Tukey’s test. F- **G**. Example traces in (**F**) at current injections of -50pA and 50pA, Scale bar=50mV, 20ms and quantification in (**G**) of evoked action potentials during incremental current injections showing decreased intrinsic excitability of CeA neurons in female SI+SNI mice (n=22 cells) as compared to female ctrl mice (n=32), Two-tailed, unpaired t- test comparing current injection eliciting half-maximal firing rates (I50), p<0.0001. **H-I**. Example traces in (**H**) and quantification in (**I**) of firing rates upon optogenetic excitation of PVT axonal terminals showing pathway-specific hyperexcitability of CeA neurons in female SI+SNI (n=20 cells) as compared to female ctrl (n=16), Two-tailed Mann-Whitney-Test, p=0.014, unpaired Cohen’s d=0.844 [95.0%CI 0.118, 1.55].

To examine whether a vulnerability-stress interaction induces specific synaptic plasticity alterations in the PVT→CeA pathway, we performed patch-clamp recordings in CeA neurons in combination with light stimulation of PVT terminals (**Fig. 2B**). Neither SI and/or SNI altered light-induced AMPAR- and NMDAR-mediated EPSC amplitudes or decay kinetics (**Fig. S5**). In order to investigate long-term plasticity and synaptic integration, we measured the ratio of AMPAR-to NMDAR-mediated EPSCs in the same CeA neuron (AMPA/NMDA ratio). We showed that neither SI nor SNI significantly changed the AMPA/NMDA ratios in male and female mice (**Fig. S5**) suggesting unaltered postsynaptic function of the PVT→CeA pathway under vulnerability-stress conditions.

To investigate whether vulnerability-stress conditions alter the presynaptic function in the PVT→CeA pathway, we stimulated PVT axons with two brief light pulses and measured the paired-pulse ratio (PPR) of AMPAR-mediated EPSCs in CeA neurons. Neither SI nor SNI altered PPRs in male mice (**Fig. 2D**). In contrast, the specific interaction of SI and SNI increased the PPR in the PVT→CeA pathway of female SI+SNI mice, in contrast to all other female groups (ctrl, SI-only, and SNI-only; **Fig. 2E**). We concluded that a vulnerability-stress interaction induces a sex-specific presynaptic facilitation in the PVT→CeA pathway of female SI+SNI mice.

Next, we investigated whether vulnerability-stress conditions increase neural excitability in CeA in comparison to ctrl mice. We applied incremental current steps (**Fig. 2F**) and observed reduced firing responses in female SI+SNI mice (**Fig. 2G**) suggesting reduced intrinsic excitability of CeA neurons under vulnerability-stress conditions. To determine whether the sex-specific short-term facilitation increases excitability of the PVT→CeA pathway under vulnerability-stress conditions in comparison to control mice, we stimulated PVT axons with brief bursts of light pulses and recorded firing responses of CeA neurons (**Fig. 2H**) in female mice. Interestingly, burst stimulation of PVT axons enhanced firing responses of CeA neurons in female SI+SNI mice (**Fig. 2I**) suggesting enhanced pathway-specific excitability in the PVT-CeA pathway under vulnerability-stress conditions.

### Vulnerability-stress conditions induce spatial hyperexcitability of the PVT→CeA pathway in anxiously avoiding mice

Based on our findings that the interaction between vulnerability and stress specifically enhances hyperalgesia, conflict-related anxiety, and short-term facilitation in the PVT-CeA pathway in female mice, we now focused only on female mice to better understand the causal mechanisms by which vulnerability-stress conditions alter female PVT→CeA circuit function and behavioral responses.

To investigate whether vulnerability-stress conditions induce a specific PVT→CeA circuit dysfunction during motivational conflict in female mice, we retrogradely expressed the genetically encoded Ca^2+^ sensor GCaMP7c (AAV-DIO-GCaMP7c) in CeA-projecting PVT neurons and implanted optical cannulas above the PVT (**Fig. 3A** and **3B**). Photometric signals from CeA-projecting PVT neurons and locomotion were recorded in female control, SI-only, SNI-only, and SI+SNI mice during the OFT and NSF (**Fig. 3C**). Calcium recordings during the OFT were used as an active control, based on our observation that vulnerability-stress conditions do not increase avoidance behavior in this test. To examine potential spatial encoding of behavioral differences between the NSF and OFT, we recorded calcium signals from CeA-projecting PVT neurons in ctrl mice. Surprisingly, our results revealed a spatially distinct activation pattern: the PVT-CeA circuit was strongly activated in the corners of the OFT (**Fig. 3D**), while in the NSF arena, it was strongly activated in the center (**Fig. 3E**) suggesting behaviorally relevant spatial coding of CeA-projecting PVT neurons.

**Figure 3.**
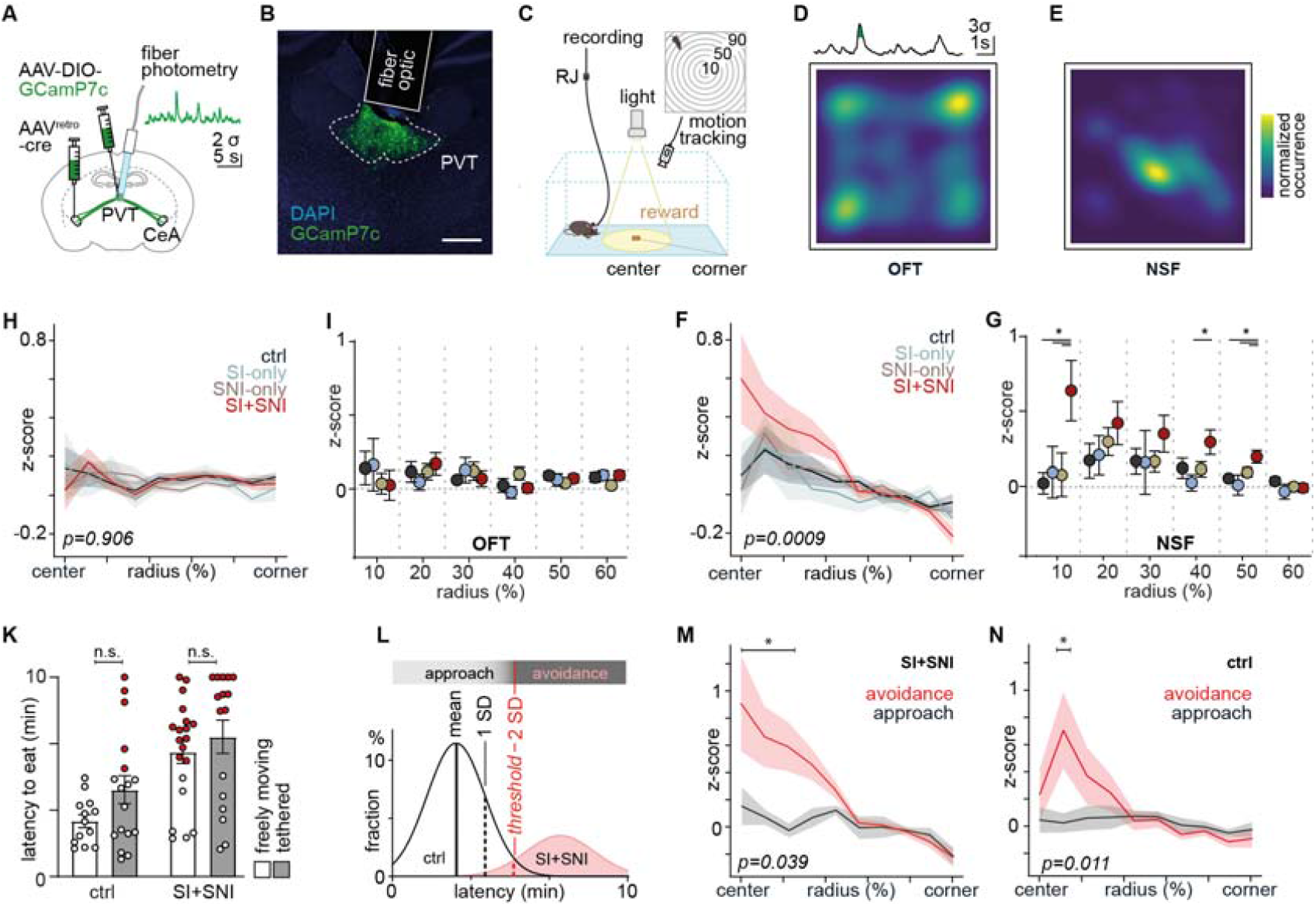
Vulnerability-stress conditions induce spatial hyperexcitability of the PVT→CeA pathway in anxiously avoiding mice A. Schematic viral strategy and fiber optic placement to record PVT→CeA pathway activity. **B**. GCamP7c expression in CeA-projecting PVT neurons, Scale bar=200µm. **C**. Schematic of fiber photometry and motion tracking during NSF test (**C**), inset shows radial gradient transformation. **D-E**. Probability density map of high calcium activity (>3 SD) in OFT (**D**) and NSF (**E**). **H-I**. Calcium activity of PVT→CeA neurons relative to the normalized radial distance of the mouse to the arena center in OFT (**H-I**) ctrl (n=16), SI-only (n=8), SNI-only (n=13) and SI+SNI (n=16) mice, F_27,414_=0.658, p(radiusXgroup)=0.906 and NSF (**F-G**) ctrl (n=17), SI-only (n=8), SNI-only (n=13), SI+SNI (n=15) mice, F_27, 434_=2.15, p(radiusXgroup)=0.0009. Two-Way ANOVA with Sidak Post test. **K**. Comparison of the latency to eat in the NSF test among freely moving and tethered mice in ctrl and SI+SNI groups. **L**. Schematic definition of an avoidance threshold at 2 SD above the mean latency to eat of untethered ctrl mice in the NSF test. Red datapoints in (**K**) represent mice above the threshold. **M-N**. Comparison of spatial calcium data of avoiding and approaching mice in the female SI+SNI (n=10 and 5) condition (**M**), F_9,117_=2.047, p(radiusXgroup)=0.0399 and female ctrl (n=5 and 12) condition (**N**), F_9,133_=2.492, p(radiusXgroup)=0.0115. Two-Way ANOVA with Sidak Post test.

To investigate whether vulnerability-stress conditions induce a spatial dysfunction in the PVT-CeA circuit specific to the NSF, we recorded average photometric activity relative to the mouse’s radial position in the OFT and NSF arena in female ctrl, SI-only, SNI-only and SI+SNI mice (**Fig. 3C** top). In OFT, female SI+SNI mice did not exhibit elevated PVT→CeA circuit activity (**Fig. 3H** and **I**) and locomotor behavior (**Fig. S6**). Remarkably, female SI+SNI mice displayed elevated calcium activity of CeA-projecting PVT neurons in the NSF center in comparison to control, SI-only and SNI-only mice (**Fig. 3F** and **G**), suggesting that vulnerability-stress conditions induce a spatial- and context-specific PVT→CeA pathway hyperexcitability during motivational conflict.

Tethering might affect behavioral flexibility during motivational conflict despite sufficient habituation. We thus compared anxious avoidance between freely moving and tethered female mice in the NSF and observed that tethering significantly affects the behavior in the NSF with non-significant increases in the latency to eat (**Fig. 3K**), suggesting that tethering heightens anxious avoidance during motivational conflict. Yet, this difference in the behavioral performance of tethered mice (**Fig. 3K**) enabled us to test whether PVT-CeA circuit dynamics depend on an anxious state. We defined an anxious avoidance threshold (latency = 309 s) as a latency increase exceeding two standard deviations (SD = 71 s) above the mean performance (mean = 167 s) of freely moving ctrl mice in the NSF test (**Fig. 3L**). Below-threshold latencies are defined as approach responses, above-threshold latencies as avoidance responses (**Fig 3L**). Interestingly, freely moving ctrl mice did not show such avoidance responses (**Fig. 3K**). Mice with an avoidance response in the NSF test showed a strong central increase in calcium activity of CeA-projecting PVT neurons (**Fig 3 M** and **N**) both in ctrl and in SI+SNI mice compared to mice with approach responses. These findings suggest that the PVT→CeA pathway encodes for state-dependent anxious avoidance under motivational conflict.

### The PVT→CeA pathway specifically mediates female hyperalgesia and anxiety in vulnerability-stress conditions

To investigate whether the PVT→CeA pathway causally mediates chronic hyperalgesia and anxious avoidance in female mice under vulnerability-stress conditions, we expressed either the inhibitory designer receptor hM4Di or the green fluorescent protein GFP for the respective control groups in the PVT→CeA pathway of female mice (**Fig. 4A**). Three weeks after surgery, all mice received intraperitoneal (i.p.) injections of the hM4Di activator clozapine-N-oxide (CNO, 3 mg/kg, i.p.) 45 min before the start of the behavioral tests (**Fig. 4B**). Interestingly, chemogenetic inhibition of the PVT→CeA pathway specifically reduced the paw withdrawal sensitivity of female SI+SNI mice in the von Frey Test (**Fig. 4D**). However, it did not affect the mechanical hyperalgesia of ctrl, SI-only and SNI-only mice, nor did it influence cold allodynia in the Cold Plate Test (**Fig. S9**) suggesting that the PVT-CeA pathway specifically mediates the additive chronic mechanical hyperalgesia under vulnerability-stress conditions, but not chronic neuropathic pain.

**Figure 4.**
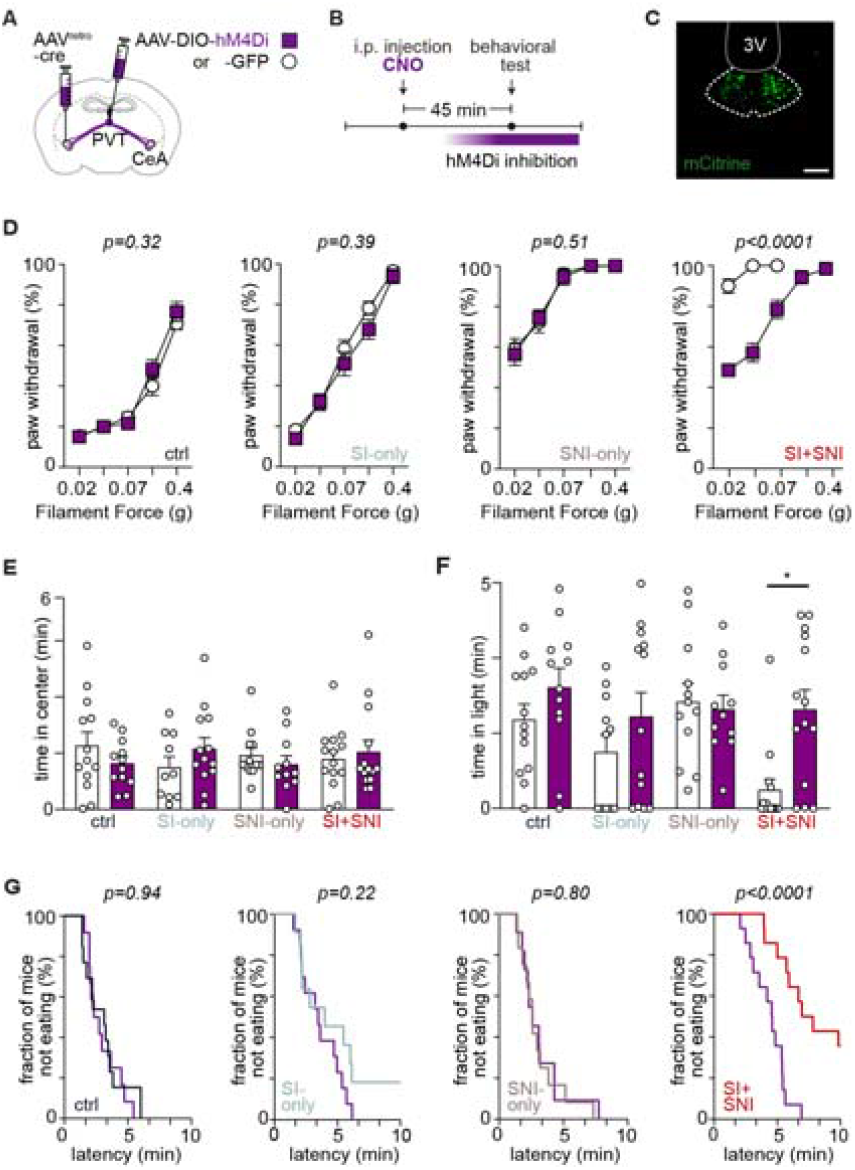
The PVT→CeA pathway selectively mediates hyperalgesia and anxiety in females under vulnerability-stress conditions. **A.** Schematic viral strategy for chemogenetic inhibition of the PVT→CeA pathway. **B**. Experimental timeline. **C**. Expression of the designer receptor hM4Di in CeA-projecting PVT neurons following viral injection. Scale bar=100 µm. **D**. Paw withdrawal responses during von Frey test measured 12 weeks post-SNI surgery. Chemogenetic inhibition of the PVT→CeA pathway reduced mechanical hypersensitivity only in female SI+SNI mice (n=14; p<0.0001). No significant effects were observed in ctrl (n=12; p>0.999), SI-only (n=13; p=0.835), or SNI-only (n=11; p=0.9108) groups compared to GFP-expressing mice, ctrl (n=13), SI-only (n=11), SNI-only (n=12), SI+SNI (n=14). **E**. Chemogenetic silencing of the PVT→CeA pathway did not affect the behavioral response in OFT. Group sizes as mentioned above. **F**. Chemogenetic silencing of the PVT→CeA pathway during LDB selectively increased the time spent in the light compartments in female SI+SNI mice (p=0.004), but not in ctrl (p=0.59), SI-only (p=0.522) and SNI-only mice (p=0.998). Group sizes as mentioned above. **G**. Effect of chemogenetic inhibition of the PVT→CeA pathway on the latency to eat in NSF, plotted in survival curves. Systemic CNO administration significantly reduced the latency to eat exclusively in female hM4Di-expressing SI+SNI mice (p<0.0001) but not in ctrl (p=0.935), SI-only (p=0.216), or SNI-only (p=0.803) groups. Log-rank (Mantel-Cox) test, Group sizes as indicated above.

To examine the role of the PVT→CeA circuit in comorbid anxious avoidance under vulnerability-stress conditions, we performed a series of anxiety-related behavioral tests. Chemogenetic inhibition of PVT→CeA pathway did not significantly affect anxiety-like behaviors in the OFT (**Fig. 4E**). However, it specifically mitigated anxious avoidance of female SI+SNI mice in LDB (**Fig. 4F**) and NSF (**Fig. 4G**). Chemogenetic inhibition did not alter behavioral responses of female ctrl, SI-only and SNI-only mice in the LDB and NSF in comparison to GFP control mice, suggesting that CeA-projecting PVT neurons may mediate anxious avoidance during motivational conflict specifically in a state of heightened anxiety, but not under baseline conditions. Additionally, chemogenetic inhibition of the PVT→CeA pathway did not affect pellet consumption during the NSF test in female SI+SNI mice, nor did it influence locomotor or exploratory behavior in the OFT across all groups (**Fig. S10)**. These results suggest that the specific chemogenetic rescue of anxious avoidance in female SI+SNI mice is not mediated by alterations in locomotor or consummatory behaviors. Overall, these results demonstrate that vulnerability-stress conditions specifically induce a hyperactive PVT→CeA circuit dysfunction in female SI+SNI mice, which causally mediates enhanced mechanical hyperalgesia and anxious avoidance.

## Discussion

Together with current knowledge on the paraventricular thalamus, our findings reveal that early adversity drives the chronification of hyperalgesia and anxious avoidance in female mice via short-term facilitation and hyperexcitability of the PVT-CeA pathway. This unknown PVT-CeA circuit dysfunction (1) depends on a history of early adversity, (2) precipitates by a subsequent nerve injury, (3) develops in a sex-specific manner in females, and (4) mediates tactile hypersensitivity in combination with anxiety - together representing key features of nociplastic primary pain conditions.

While few studies have examined chronic pain and comorbid anxiety in a vulnerability-stress framework in mice^28^, Nishinaka et al. showed that combining maternal separation (days 15-21) with early social isolation (from day 21) resulted in pronounced behavioral deficits. These included anxiety-like avoidance in female mice in the OFT and thermal and tactile hypersensitivity in both sexes for up to two weeks after nerve injury. We attenuated this vulnerability-stress model by focusing on the clinically relevant period of peripubertal adversity^29^ using early social isolation alone and investigated mice for up to 12 weeks after nerve injury. The overall similar but less pronounced behavioral deficits in our model may be due to the absence of an additional prepubertal adversity, which may affect male mice differently, or due to the longer follow-up period in our study, which addresses a chronic disease state. Interestingly, vulnerability-stress conditions did not specifically increase conflict anxiety or hyperalgesia in male mice. This observed sex difference may be due to protective endocrine or social factors, unique neurobiological alterations to early adversity or alternative coping strategies that lead to other behavioral deficits in male mice. Our attenuated vulnerability-stress model thus provides a suitable animal model for the translational investigation of sex-specific trajectories of different mental deficits after early adversity.

Avoidance has been implicated as a cardinal symptom of anxiety disorders and is thought to be an underlying mechanism that perpetuates anxiety^30^. The balance between approach and avoidance tendencies during motivational conflict, like in Pavlovian counterconditioning^31,32^, is regulated via a dynamic interplay between opposing activities of PVT subsets^16,33^. Our results unravel unknown comparable dynamics in the PVT-CeA circuit that mediate avoidance behavior under vulnerability-stress conditions. However, this dynamic over-excitation of the PVT-CeA circuit occurs in a state-dependent manner only in anxiously avoiding mice and thus may track strong anxiety. In contrast, other PVT projections, e.g. to nucleus accumbens, may regulate approach tendencies during motivational conflict in control mice^33^. A state of heightened anxiety in female mice under vulnerability-stress conditions may reduce behavioral flexibility via recurrent dynamic activation of CeA-projecting PVT(D2-) neurons driving pursuit avoidance^33^. However, the neural origin of this PVT-CeA circuit dysfunction remains unknown. Early social isolation induces sociability deficits via reduced drive of PVT-projecting neurons of the prelimbic cortex (PrL). As PVT, suggesting that other afferent inputs mediate PVT-CeA circuit dysfunction under vulnerability-stress conditions. A potential may arise from increased dopaminergic PVT disinhibition by the locus coeruleus^34^, which mediates stress responsiveness.

The PVT-CeA circuit also mediates the establishment and long-term maintenance of conditioned fear associations^5,10,16-17^ via a presynaptic depression in SOM+ CeA neurons and a presynaptic facilitation in SOM-CeA neurons^35^. Vulnerability-stress conditions - similar to fear conditioning^35^ - induce presynaptic facilitation in the PVT-CeA circuit, which can be explained as activity-dependent short-term plasticity upon recurrent circuit excitation. A subsequent reduction of the initial glutamate release probability or increased residual calcium^36^ may thus increase the pathway-specific excitability of CeA neurons when PVT afferents fire in longer bursts. The parallel reduction of intrinsic excitability under vulnerability-stress conditions may enable CeA neurons to maintain homeostasis despite increased input-specific drive^27^. Although an excitation of the PVT-CeA circuit drives mechanical hyperalgesia in rodents, it only mediates subacute neuropathic pain up to two weeks after nerve lesion but not more chronic hyperalgesia in male mice^18,37^. Our findings now reveal a previously unknown role for the PVT-CeA pathway in exacerbating chronic neuropathic pain in female mice under vulnerability-stress conditions. The finding that early-life stress does not impact thermal hyperalgesia may be due to differences in specific pain modalities or to an anxious microstate during the von Frey test^38^ that allows anticipation of subsequent pain stimuli. Furthermore, the PVT-CeA circuit specifically mediates an additive mechanical hyperalgesia on top of neuropathic pain, but not the neuropathic pain itself - similar to a bottom-up subtype of nociplastic pain^39^.

Despite these findings, our study is limited by the choice of one specific vulnerability-stress model. Given the diversity of childhood adversities in patients, further research is needed to determine whether different early-life stressors result in similar PVT-CeA circuit dysfunction. Such experiments could help define a vulnerable time window during childhood. Future studies should also elucidate the neural or endocrine origin and human correlates of the PVT-CeA circuit dysfunction in female patients with chronic primary pain which would broaden the implications and foster the development of more targeted therapeutic interventions for chronic pain patients.

The results of this study, when taken together with current knowledge, particularly support the PVT-CeA pathway as a integrative hub whose adaptive modification through early adversity drives chronic disease manifestation in adulthood; our findings thus integrate the theory of vulnerability for mental disorders and the fear-avoidance model of chronic pain with a novel biological mechanism. Understanding such a neural mechanism that facilitates chronic pain holds promise to ultimately reverse the pervasive consequences of childhood adversity.

## Acknowledgments

We thank Ross Folkard for his valuable feedback on the manuscript, Rick Bernardi and Elisabeth Roebel for their technical assistance, and Aya Elsify for her support with graphical design. We thank Amit Agarwal, Christoph Körber and Anke Tappe Theodor for their help in establishing our methods. We would like to thank Jonas Tesarz for his valuable input planning and discussing the project. We are also grateful to all members of the CRC1158 community for their support and the insightful discussions on our data.

## Author contributions

Conceptualization: R.S., S.W..; Formal analysis: A.M.B., T.K., L.R., M.V.R., N.L., S.W.; Funding acquisition: R.S., S.W..; Investigation: A.M.B., T.K., L.R., M.V.R., C.C-L., S.W.; Methodology: M.V.R., N.L., L.B.; Project administration: A.M.B., S.W., R.S.; Resources: H.C.F., T.K., R.K., R.S., S.W.; Software: N.L., L.B., M.V.R.; Supervision: R.S., S.W.; Validation: S.W.; Visualization: A.M.B., S.W.; Writing – original draft: S.W.; Writing – review & editing: A.M.B., T.K., L.R., M.V.R., C.C-L., N.L., T.K., R.K., R.S., S.W.

## Methods

### Animals

Male and female C57BL6/N mice (Mus musculus) were used for all experiments. with food and water ad libitum under a 12 h :12 h light-dark cycle (light off at 7:00 am). All the experimental procedures were approved by the local governing body (Regierungspräsidium Karlsruhe, Germany, Ref. 35-9185.85/G-307/19, 35-9185.81/G-208/15 and 35-9185.81/G-247/21) and adhered to the ethical guidelines of the University Heidelberg. The experimenters were blinded to the identity of the mice during the behavioral tests and immunohistochemical quantification. ARRIVE guidelines were followed.

### Vulnerability-Stress Model

In humans, childhood adversity is often social adversity, whereas acute stressors can be of various (often non-social) kinds. In our translational mouse model, we thus investigated the interaction of social isolation (as a diathesis) and a spared nerve injury (as a somatic stressor). At postnatal day 21, male and female mice were randomly assigned to either group-housing (two to four mice per cage) or social isolation (SI, one mouse per cage) for the whole experimental period. Between week 5 and 7, male or female SI or group-housed mice were randomly assigned to receive either a spared-nerve injury (inducing neuropathic pain) or a sham surgery. Male and female mice were thus investigated in 4 experimental groups: sham group (group-housed, sham surgery); SI-only group (SI; sham surgery), SNI-only group (group-housed, SNI surgery) and SI+SNI group. As a chronic disease state, we examined the respective mice 6-12 weeks after surgery.

### Spared nerve injury (SNI)

As described previously^40^, mice were anesthetized with isoflurane anesthesia (induction: 4%; maintenance: 1.5 - 2.5%). An incision lateral to the skin surface of the right shaved thigh was made to dissect the biceps femoris muscle and expose the sciatic nerve and the branches. The tibial and the common peroneal nerves were ligated firmly then a section of the nerve bundle was removed from the ligation. The sural nerve was kept safely intact during the surgery. In sham operations, the sciatic nerve and its three branches were exposed but left intact. The muscle was repositioned and the incision was sutured subsequently. Mice were placed on the warm plate for recovery.

### Brain virus injection and fiber optic implantation

All the AAV virions employed in this study were purchased from Addgene company and the viral vector facility of Zurich University, Switzerland. Details of the virions can be seen in Table S2. Virus injection was performed via borosilicate glass capillary (Drummond, Broomall, PA, USA) using an oil hydraulic microinjector (Narishige, Tokyo, Japan). Injection and implantation coordinates can be found in Table S3. The coordinates for all the brain virus injections and fiber implantations were referred to the mouse atlas Paxinos and Franklin and are shown with respect to bregma (anterior-posterior; mediolateral) or brain surface (depth). Additional information on anatomical aspects was derived from resources of the Allen Institute for Brain Science (ALLEN Mouse Brain Atlas – Version 2; available from: http://atlas.brain-map.org/). Two injection positions were employed to thoroughly cover the posterior part of the PVT. 200nl virus solution was injected in the PVT at -1.3mm and -1.6mm anterior-posterior, at -0.5mm mediolateral and at -2.9mm depth with a 10^°^ angle towards the midline. For CeA injections 250 nl virus solution was injected per side at -1.3mm anterior-posterior, at +/-2.85mm mediolateral and at -4.9mm depth. For pathway-specific expression in the PVT→CeA pathway, AAVrg-Syn-Cre or AAVrg-Syn-Cre-mCherry virions were injected in the CeA. For in-vitro optogenetic experiments, AAV-Syn-ChR2-EYFP virions were injected in the PVT. For chemogenetic experiments, AAV-Syn-DIO-hM4Di-mCitrine or the corresponding control virions AAV-Syn-EGFP were injected in the PVT. In fiber photometric experiments, AAV-Syn-DIO-GCamP7c virions were injected in PVT. For in-vivo optogenetic stimulation (Thorlabs: 400 µm, 0.39 NA) and for fiber photometry (DORIC: 400 µm, 0.66 NA) we implanted one fiber optic cannula around 0.3 mm above the PVT at -1.5mm anterior-posterior, -0.5mm mediolateral and at -2.6mm depth with a 10^°^ angle towards the midline. The optic cannulas were fixed with dental cement (CB medical, Superbond, Japan). All the mice were kept at least 3 weeks after the brain injection for thorough virus expression and recovery. Correct virus expression and fiber optic position was confirmed on each mouse at the end of the experiments or otherwisely excluded.

### Viral vectors

For all experiments, commercially available adeno-associated virus (AAV) particles were procured from Addgene. For the DREADD experiments, the AAV vectors utilized were as follows: hM4Di encoding vector (AAV8-hSyn-DIO-HA-hM4Di-IRES-mCitrine, concentration: 1×10^^^13 viral genomes (vg)/ml, Addgene Cat#50455), retrograde virus (AAVrg-Ef1a-mCherry-IRES-Cre, concentration: 7×10^^^12 vg/ml, Addgene Cat#55632), and eGFP encoding vector (AAV8-hSyn-DIO-EGFP, concentration: 1×10^13 vg/ml, Addgene Cat#50457). These vectors were employed to selectively modulate the inhibition of PVT projections to the CeA. For electrophysiological experiments we used constitutive ChR2-eYFP encoding vector (AAV1-hSyn-hChR2(H134R)-EYF, concentration: 1×10^13^ vg/mL, Addgene Cat#26973). All viruses were aliquoted into 10 μL aliquots, stored at −80^°^ C, and thawed immediately before injection.

### Behavioral Tests

All the behavioral tests were conducted in the dark cycle of the light-dark rhythm. For chemogenetic manipulations, mice were administered with clozapine-N-oxide (CNO, 3 mg/kg, i.p.; Tocris, USA,) 45 min prior to each test. Mice for von Frey measurements were acclimatized 1 h in the test setups twice per day and three days in advance.

#### von Frey measurements

Each mouse was given a period of acclimatization within small plastic boxes (7.5□×□7.5□×□15 cm) on a wire grid floor. This acclimatization phase lasted for a minimum of 30 minutes before any testing commenced. Mechanical sensitivity of the right hindpaw was tested with manual repeated applications of von Frey filaments to the plantar surface in the order of ascending forces (0.02, 0.04, 0.07, 0.16, 0.40 g, 5 applications per filament, 30 s interval).

#### Cold plate test

Cold sensitivity was tested by measuring the withdrawal latency of the mice hindpaws on a plate maintained at a constant temperature of 2^°^ C ± 1^°^ C. A 30 s cut-off was set to prevent potential injury on the paws.

#### Open field test

Mice were placed in a 40 × 40 cm arena to freely explore and were recorded for 20 min with dim illumination (30 lux) during the test. Locomotion parameters, such as difference in distance travelled during the test and the center time, were analyzed via the EzTrack software (REF) and by Any-Maze video tracking system (Version 7.0, Stoelting, USA).

#### Light-Dark box test

The Light-Dark Box consisted of an open-topped rectangular Plexiglas box measuring 45 cm × 30 cm × 30 cm. The box was partitioned into a small area (18 cm × 30 cm) and a large area (27 cm × 30 cm) connected by a door (7.5 cm × 7.5 cm) positioned at floor level in the center of the partition. The small compartment was painted black and maintained in darkness, while the large compartment was painted white and brightly illuminated (400 lux). Each trial started in the dark compartment and lasted 5 minutes. The time spent in the light compartment, latency to reach the light compartment, and the number of transitions was recorded using the Any-Maze video tracking system (version 7.0, Stoelting, USA).

#### Novelty Suppressed Feeding (NSF) test

The Novelty-Suppressed Feeding (NSF) test is designed to induce a conflict between the innate drive to eat and the fear of approaching into the center of a bright, novel arena. The NSF arena consists of a plastic box measuring 40 x 40 x 30 cm illuminated with a 800 lux lamp and a floor covered with 2 cm of clean bedding. A single food pellet is placed in the center of the arena. Mice are transferred to the testing room at least 1 hour before testing. After 24 hours of food restriction, each mouse is placed in one corner of the testing arena. During a 10 min time period, the latency to bite into the food pellet was recorded. Mice that do not consume the food within the designated 10 min period are assigned a latency of 600 seconds. Immediately following the NSF test, mice are transferred back to their home cages and provided access to a single food pellet for 5 minutes. Latency to eat and the amount of pellet consumed in the home cage serve as control behaviors for comparison with the NSF test results.

### Slice electrophysiology

Coronal CeA slices (250 μm) were prepared using a vibratome (Leica VT1200S) and ice-cold N-methyl-D-glucamine (NMDG)-based cutting solution (135 mM NMDG, 1 mM KCl, 1.2 mM KH2PO4, 1.5 mM MgCl2, 0.5 mM CaCl2, 12.95 mM D-Glucose, 20 mM Choline bicarbonate, titrated to pH 7,4 with HCl), saturated with carbogen (95% O2 and 5% CO2). Sections that included CeA region were captured and stored in carbogenated artificial cerebrospinal fluid (ACSF; 125 mM NaCl, 2.5 mM KCl, 1.25 mM NaH2PO4, 25 mM NaHCO3, 25 mM D-Glucose, 1 mM MgCl2, 2 mM CaCl2) at 37°C for 30 min for stabilization, later at room temperature. For voltage-clamp recordings, pipettes were filled with a Cesium-based internal solution (105mM Cs-Gluconate, 25mM CsCl, 10mM Hepes, 10mM Phosphocreatine, 4mM ATP-Mg-salt, 0.3mM Guanosintriphosphate, 2.5mM QX-314-chloride, titrated with CsOH (pH 7.3). For current-clamp recordings pipettes were filled with Potassium-based internal solution (120mM K-Gluconate, 15mM KCl, 10mM Hepes, 7mM Phosphocreatine, 4mM ATP-Mg-salt, 4mM MgCl2, 0.3mM Guanosintriphosphate, 0.1mM EGTA (KOH), buffered with KOH (pH 7.2). 0.2% Biocytin was freshly added to the internal solution for histological localization of the cell. Patch-clamp recordings were only performed after correct viral expression in PVT was visually controlled before the start of every experiment. Patch pipettes were pulled from borosilicate glass (World Precision Instruments, Sarasota, Florida, USA) using a Micropipette Puller Model P-97 (Sutter Instruments, Novato, CA, USA) and had open tip resistances of 4-6 MΩ. Patch-clamp recordings from CeA neurons were performed using an EPC-10 Double USB amplifier controlled by PatchMaster software (HEKA, Lambrecht, Germany). Recorded current and voltage traces were digitized at 200 kHz and Bessel-filtered (2.9 kHz). Optogenetic stimulation of PVT terminals in the CeA was performed using a fiber-coupled LED (M470F3, Thorlabs). Brief light pulses (5 ms) were delivered via an optic fiber (400μm, M118L02, Thorlabs) placed ∼1-2 mm away from the recorded cell. All whole cell recordings were performed in presence of 100 μM Picrotoxin (HelloBio, Bristol, UK) to block GABAergic transmission. AMPA currents were recorded at a membrane potential of -70mV, NMDA currents at +40mV after 10 μM CNQX (HelloBio, Bristol, UK) was added. NMDA currents were blocked by 50 μM DL-AP5 (HelloBio, Bristol, UK). EPSC properties were determined from an average of 20 EPSCs per cell using IGOR (Wavemetrics, Lake Oswego, OR) with TaroTools and NeuroMatic extensions. Unstable recordings in which access resistance exceeded 20 MΩ or varied more than 20% over time were excluded. AMPAR/NMDAR ratios were calculated by dividing the amplitude of the AMPAR-mediated EPSC by the amplitude of the NMDAR-mediated EPSC. The paired-pulse ratio was recorded using two light pulses at 10 Hz frequency and calculated by dividing the amplitude of the second EPSC by the first EPSC.

### In-vivo fiber photometric recordings

Fiber photometry was performed as previously described^16^. Briefly, mice were allowed to habituate to the fiber patch cord for approximately 5 min before each behavior test. GCaMP7c fluorescence and isosbestic autofluorescence signals were excited by the fiber photometry system (Doric Lenses) using two sinusoidally modulated 473 nm (211 Hz) and 405 nm (531 Hz) LEDs (DC4100, ThorLabs). Both LEDs were combined via a commercial Mini Cube fiber photometry apparatus (Doric Lenses) into a single fiber patch cord (400-μm core, 0.57 NA, Doric Lenses) connected to the brain implant in each mouse. The light intensity at the interface between the fiber tip and the animal was adjusted from 10 μW to 20 μW and remained constant throughout each session. A lock-in amplifier (Photometry Console, Doric Lenses) with software (Neuroscience Studio V5, Doric Lenses) recorded and saved real-time demodulated emission signals and behavior-relevant TTL inputs (filament touch, air puff, electric shock, water reward).. During anxiety tests, the behavior of an individual mouse inside the arena was tracked with synchronized camera recordings (The Imaging Source, DMK33UX287) that were post-hoc analyzed using ezTrack (Pennington, 2019).

### Histology

For histological analysis, animals were deeply anesthetized with pentobarbital (100-150mg/kg) and then transcardially perfused with phosphate-buffered saline (PBS, pH 7.4, 4□^°^C), followed by a paraformaldehyde (PFA 4% in PBS, 4□°C) solution. After extraction, the brains were post-fixed in 4% PFA at 4□ ^°^C for 24 hours. Coronal brain slices (60 µm) were cut using a vibratome (Leica, VT1000S). For experiments in Fig 4, brains were cryoprotected by immersion in a 30% PBS-buffered sucrose solution until saturation was achieved (for over 24 hours). Coronal brain sections measuring 20 μm were cut using a freezing microtome (SM 2010R, Leica). For reconstruction of optical cannula position, brains (including implants) were kept in 4% PFA for 14 days, before washed in PBS.

### Data Analysis

#### Preprocessing

Calcium data was analysed using the Python programming language. Preprocessing of the raw data was performed as previously described (Martianova 2019). In brief, for each trial, after smoothing of the data with a moving window average, baseline drift correction using the adaptive iteratively reweighted penalized least squares (airPLS) algorithm was performed (Zhang, 2010). Subsequently, autofluorescence signals (F405nm) were subtracted from GCamp7 signals (F473nm) to control for movement artifacts. To achieve intersession and intersubject comparability, zdF/F was calculated as the standard z-score of one session using the formula zdF/F = signal – mean(signal) / SD(signal) (Sherathiya, 2021). Unless otherwise specified the numpy library was used for all calculations.

#### NSF and OF

In order to correlate spatial information with calcium dynamics the location of the mouse obtained from video analysis was transformed and represented as the normalized radial distance from the centre for each session. After binning to 10 equally sized segments the mean calcium data of each mouse was averaged to obtain the group average and calculate the standard error of the mean. Using the ‘curve fit’ function of the scipy library a linear fit was performed and fit parameters extracted. Correlation between groups was determined using the numpy ‘corrcoef’ function. All plots were created using the matplotlib library.

### Statistics

Unless specified otherwise, data is shown as mean ± standard error of the mean (SEM). For comparisons between two groups, paired or unpaired two-tailed t-tests were used for dependent and independent measurements respectively. In cases where no Gaussian distribution could be assumed, the non-parametric Mann-Whitney test was used. For comparison between three or more groups one-way Anova was used, followed by Bonferroni’s post hoc test for multiple comparisons. For comparison of data influenced by two different variables or repeated measures two-way Anova was used, followed by Tukey’s and Sidak’s post-hoc comparisons. In cases where no Gaussian distribution could be assumed, the non-parametric Kruskal-Wallis test with Dunn’s multiple comparisons was used. Behaviour analysis of the NSF-test was done using Kaplan-Meier survival curves with log-rank (Mantel-Cox) test and Bonferroni correction to determine statistical significance. Where effect sizes are indicated, Cohen’s d was used. The level of significance was set to 0.05 across all data analyses. GraphPad Prism (Version 6) and the Scipy Python library (Version 1.10.0) were used for statistical analysis.

## References

1. Vlaeyen, J.W.S., and Linton, S.J. (2000). Fear-avoidance and its consequences in chronic musculoskeletal pain: a state of the art. PAIN 85, 317. 10.1016/S0304-3959(99)00242-0.

2. Bushnell, M.C., Čeko, M., and Low, L.A. (2013). Cognitive and emotional control of pain and its disruption in chronic pain. Nat. Rev. Neurosci. 14, 502–511. 10.1038/nrn3516.

3. Zubin, J., and Spring, B. (1977). Vulnerability--a new view of schizophrenia. J. Abnorm. Psychol. 86, 103–126. 10.1037//0021-843x.86.2.103.

4. Yamada, K., Matsudaira, K., Tanaka, E., Oka, H., Katsuhira, J., and Iso, H. (2017). Sex-specific impact of early-life adversity on chronic pain: A large population-based study in japan. J. Pain Res. 10, 427–433. 10.2147/JPR.S125556.

5. Heim, C., and Nemeroff, C.B. (2001). The role of childhood trauma in the neurobiology of mood and anxiety disorders: preclinical and clinical studies. Biol. Psychiatry 49, 1023–1039. 10.1016/S0006-3223(01)01157-X.

6. McLean, C.P., Asnaani, A., Litz, B.T., and Hofmann, S.G. (2011). Gender differences in anxiety disorders: prevalence, course of illness, comorbidity and burden of illness. J. Psychiatr. Res. 45, 1027–1035. 10.1016/j.jpsychires.2011.03.006.

7. Tolin, D.F., and Foa, E.B. (2006). Sex differences in trauma and posttraumatic stress disorder: A quantitative review of 25 years of research. Psychol. Bull. 132, 959–992. 10.1037/0033-2909.132.6.959.

8. Goodwill, H.L., Manzano-Nieves, G., Gallo, M., Lee, H.-I., Oyerinde, E., Serre, T., and Bath, K.G. (2019). Early life stress leads to sex differences in development of depressive-like outcomes in a mouse model. Neuropsychopharmacol. Off. Publ. Am. Coll. Neuropsychopharmacol. 44, 711–720. 10.1038/s41386-018-0195-5.

9. Bondar, N.P., Lepeshko, A.A., and Reshetnikov, V.V. (2018). Effects of Early-Life Stress on Social and Anxiety-Like Behaviors in Adult Mice: Sex-Specific Effects. Behav. Neurol. 2018, 1538931. 10.1155/2018/1538931.

10. Candemir, E., Post, A., Dischinger, U.S., Palme, R., Slattery, D.A., O’Leary, A., and Reif, A. (2019). Limited effects of early life manipulations on sex-specific gene expression and behavior in adulthood. Behav. Brain Res. 369, 111927. 10.1016/j.bbr.2019.111927.

11. P Graf, A., Hansson, A.C., and Spanagel, R. (2024). Isolated during adolescence: long-term impact on social behavior, pain sensitivity, and the oxytocin system in male and female rats. Biol. Sex Differ. 15, 78. 10.1186/s13293-024-00655-7.

12. Penzo, M.A., Robert, V., Tucciarone, J., De Bundel, D., Wang, M., Van Aelst, L., Darvas, M., Parada, L.F., Palmiter, R.D., He, M., et al. (2015). The paraventricular thalamus controls a central amygdala fear circuit. Nature 519, 455–459. 10.1038/nature13978.

13. Kooiker, C.L., Chen, Y., Birnie, M.T., and Baram, T.Z. (2023). Genetic Tagging Uncovers a Robust, Selective Activation of the Thalamic Paraventricular Nucleus by Adverse Experiences Early in Life. Biol. Psychiatry Glob. Open Sci. 10.1016/j.bpsgos.2023.01.002.

14. Zhu, Y., Nachtrab, G., Keyes, P.C., Allen, W.E., Luo, L., and Chen, X. (2018). Dynamic salience processing in paraventricular thalamus gates associative learning. Science 362, 423–429. 10.1126/science.aat0481.

15. McNally, G.P. (2021). Motivational competition and the paraventricular thalamus. Neurosci. Biobehav. Rev. 125, 193–207. 10.1016/j.neubiorev.2021.02.021.

16. Ma, J., du Hoffmann, J., Kindel, M., Beas, B.S., Chudasama, Y., and Penzo, M.A. (2021). Divergent projections of the paraventricular nucleus of the thalamus mediate the selection of passive and active defensive behaviors. Nat. Neurosci. 24, 1429–1440. 10.1038/s41593-021-00912-7.

17. Do-Monte, F.H., Quiñones-Laracuente, K., and Quirk, G.J. (2015). A temporal shift in the circuits mediating retrieval of fear memory. Nature 519, 460–463. 10.1038/nature14030.

18. Liang, S.-H., Zhao, W.-J., Yin, J.-B., Chen, Y.-B., Li, J.-N., Feng, B., Lu, Y.-C., Wang, J., Dong, Y.-L., and Li, Y.-Q. (2020). A Neural Circuit from Thalamic Paraventricular Nucleus to Central Amygdala for the Facilitation of Neuropathic Pain. J. Neurosci., JN-RM-2487-19. 10.1523/JNEUROSCI.2487-19.2020.

19. Chang, Y.-T., Chen, W.-H., Shih, H.-C., Min, M.-Y., Shyu, B.-C., and Chen, C.-C. (2019). Anterior nucleus of paraventricular thalamus mediates chronic mechanical hyperalgesia. Pain 160, 1208–1223. 10.1097/j.pain.0000000000001497.

20. Chen, W.-K., Liu, I.Y., Chang, Y.-T., Chen, Y.-C., Chen, C.-C., Yen, C.-T., Shin, H.-S., and Chen, C.-C. (2010). Ca(v)3.2 T-type Ca2+ channel-dependent activation of ERK in paraventricular thalamus modulates acid-induced chronic muscle pain. J. Neurosci. Off. J. Soc. Neurosci. 30, 10360–10368. 10.1523/JNEUROSCI.1041-10.2010.

21. Sharp, T.J., and Harvey, A.G. (2001). Chronic pain and posttraumatic stress disorder: mutual maintenance? Clin. Psychol. Rev. 21, 857–877. 10.1016/S0272-7358(00)00071-4.

22. Jensen, K.B., Kaptchuk, T.J., Chen, X., Kirsch, I., Ingvar, M., Gollub, R.L., and Kong, J. (2015). A Neural Mechanism for Nonconscious Activation of Conditioned Placebo and Nocebo Responses. Cereb. Cortex 25, 3903–3910. 10.1093/cercor/bhu275.

23. Kirouac, G.J. (2021). The Paraventricular Nucleus of the Thalamus as an Integrating and Relay Node in the Brain Anxiety Network. Front. Behav. Neurosci. 15, 627633. 10.3389/fnbeh.2021.627633.

24. Kooiker, C.L., Birnie, M.T., and Baram, T.Z. (2021). The Paraventricular Thalamus: A Potential Sensor and Integrator of Emotionally Salient Early-Life Experiences. Front. Behav. Neurosci. 15.

25. Pliota, P., Böhm, V., Grössl, F., Griessner, J., Valenti, O., Kraitsy, K., Kaczanowska, J., Pasieka, M., Lendl, T., Deussing, J.M., et al. (2020). Stress peptides sensitize fear circuitry to promote passive coping. Mol. Psychiatry 25, 428–441. 10.1038/s41380-018-0089-2.

26. Teicher, M.H., Samson, J.A., Anderson, C.M., and Ohashi, K. (2016). The effects of childhood maltreatment on brain structure, function and connectivity. Nat. Rev. Neurosci. 17, 652–666. 10.1038/nrn.2016.111.

27. Samuels, B.A., and Hen, R. (2011). Novelty-Suppressed Feeding in the Mouse. In Mood and Anxiety Related Phenotypes in Mice: Characterization Using Behavioral Tests, Volume II Neuromethods., T. D. Gould, ed. (Humana Press), pp. 107–121. 10.1007/978-1-61779-313-4_7.

28. Nishinaka, T., Nakamoto, K., and Tokuyama, S. (2015). Enhancement of nerve-injury-induced thermal and mechanical hypersensitivity in adult male and female mice following early life stress. Life Sci. 121, 28–34. 10.1016/j.lfs.2014.11.012.

29. Hodes, G.E., and Epperson, C.N. (2019). Sex Differences in Vulnerability and Resilience to Stress Across the Life Span. Biol. Psychiatry 86, 421–432. 10.1016/j.biopsych.2019.04.028.

30. Robin L. A.,, and Martin, P.P. (2010). Neural systems underlying approach and avoidance in anxiety disorders. Dialogues Clin. Neurosci. 12, 517–531.

31. Choi, E.A., Jean-Richard-Dit-Bressel, P., Clifford, C.W.G., and McNally, G.P. (2019). Paraventricular Thalamus Controls Behavior during Motivational Conflict. J. Neurosci. Off. J. Soc. Neurosci. 39, 4945–4958. 10.1523/JNEUROSCI.2480-18.2019.

32. Choi, E.A., and McNally, G.P. (2017). Paraventricular Thalamus Balances Danger and Reward. J. Neurosci. Off. J. Soc. Neurosci. 37, 3018–3029. 10.1523/JNEUROSCI.3320-16.2017.

33. Beas, S., Khan, I., Gao, C., Loewinger, G., Macdonald, E., Bashford, A., Rodriguez-Gonzalez, S., Pereira, F., and Penzo, M.A. (2024). Dissociable encoding of motivated behavior by parallel thalamo-striatal projections. Curr. Biol. CB 34, 1549-1560.e3. 10.1016/j.cub.2024.02.037.

34. Beas, B.S., Wright, B.J., Skirzewski, M., Leng, Y., Hyun, J.H., Koita, O., Ringelberg, N., Kwon, H.-B., Buonanno, A., and Penzo, M.A. (2018). The locus coeruleus drives disinhibition in the midline thalamus via a dopaminergic mechanism. Nat. Neurosci. 21, 963–973. 10.1038/s41593-018-0167-4.

35. Li, H., Penzo, M.A., Taniguchi, H., Kopec, C.D., Huang, Z.J., and Li, B. (2013). Experience-dependent modification of a central amygdala fear circuit. Nat. Neurosci. 16, 332–339. 10.1038/nn.3322.

36. Hanse, E., and Gustafsson, B. (2001). Paired-Pulse Plasticity at the Single Release Site Level: An Experimental and Computational Study. J. Neurosci. 21, 8362–8369. 10.1523/JNEUROSCI.21-21-08362.2001.

37. Tang, Q.-Q., Wu, Y., Tao, Q., Shen, Y., An, X., Liu, D., and Xu, Z. (2024). Direct paraventricular thalamus-basolateral amygdala circuit modulates neuropathic pain and emotional anxiety. Neuropsychopharmacology 49, 455–466. 10.1038/s41386-023-01748-4.

38. Signoret-Genest, J., Schukraft, N. L, Reis, S., Segebarth, D., Deisseroth, K., and Tovote, P. (2023). Integrated cardio-behavioral responses to threat define defensive states. Nat. Neurosci. 26, 447–457. 10.1038/s41593-022-01252-w.

39. Kaplan, C.M., Kelleher, E., Irani, A., Schrepf, A., Clauw, D.J., and Harte, S.E. (2024). Deciphering nociplastic pain: clinical features, risk factors and potential mechanisms. Nat. Rev. Neurol. 20, 347–363. 10.1038/s41582-024-00966-8.

40. Decosterd, I., and Woolf, C.J. (2000). Spared nerve injury: an animal model of persistent peripheral neuropathic pain. Pain 87, 149–158. 10.1016/S0304-3959(00)00276-1.

